# The Greenland shark (*Somniosus microcephalus*) genome provides insights into extreme longevity

**DOI:** 10.1101/2024.09.09.611499

**Authors:** Arne Sahm, Alexander Cherkasov, Hequn Liu, Danila Voronov, Kanstantsin Siniuk, Robert Schwarz, Oliver Ohlenschläger, Silke Förste, Martin Bens, Marco Groth, Ivonne Görlich, Sonia Paturej, Sven Klages, Bjoern Braendl, Jesper Olsen, Peter Bushnell, Amalie Bech Poulsen, Sara Ferrando, Fulvio Garibaldi, Davide Lorenzo Drago, Eva Terzibasi Tozzini, Franz-Josef Müller, Martin Fischer, Helene Kretzmer, Paolo Domenici, John Fleng Steffensen, Alessandro Cellerino, Steve Hoffmann

## Abstract

The Greenland shark (*Somniosus microcephalus*) is the longest-lived vertebrate known, with an estimated lifespan of ∼ 400 years. Here, we present a chromosome-level assembly of the 6.45 Gb Greenland shark, rendering it one of the largest non-tetrapod genomes sequenced so far. Expansion of the genome is mostly accounted for by a substantial expansion of transposable elements. Using public shark genomes as a comparison, we found that genes specifically duplicated in the Greenland shark form a functionally connected network enriched for DNA repair function. Furthermore, we identified a unique insertion in the conserved C-terminal region of the key tumor suppressor p53. We also provide a public browser to explore its genome.

## Introduction

Life-history traits show significant variation in vertebrates, and the evolution of life-history traits is profoundly shaped by adaptations to habitat and ecological conditions. Lifespan shows robust covariation with adult size, metabolism, and age at sexual maturity across different taxa (Speakman 2005; Magalhães et al. 2007; Ricklefs 2010). Yet, long-lived outliers exist, e.g., Galapagos giant tortoise, bowhead whale, clownfish, rockfish, naked mole-rat, or Brandt’s bat (Holtze et al. 2021). The term “negligible senescence” has been coined for such long-lived animals that appear to escape age-dependent deterioration (Finch 1998). However, only limited support has been available for this hypothesis. It has been proposed that an overarching principle in the evolution of exceptional longevity is the occupancy of habitats that are not easily accessible. For instance, long-living species are found among troglomorph (Voituron et al. 2011) and hypogean (Edrey et al. 2011) taxa. In marine environments, depth may be associated with longevity, as suggested by the exceptional (> 100 y) lifespan of the rockfish (genus Sebastes) (Cailliet et al. 2001; Kolora et al. 2021). Consequently, molecular studies focusing on interspecies comparisons have been initiated to unravel the mechanisms and genetic architecture of life-history variation and identified signaling pathways that show signatures of selection in multiple clades that evolved exceptional longevity. Longevity-associated genes mainly relate to DNA repair, mitochondrial balance/mitochondrial biogenesis, immune system functions, oxidative stress resistance, and telomere maintenance (Holtze et al. 2021). For example, in Sebastes, copy number expansions in the immune modulatory butyrophilin gene family are associated with longevity (Kolora et al. 2021).

The Greenland shark (GLS, *Somniosus microcephalus*) is a giant (>5 m) iconic abyssal species that inhabits the North Atlantic depths and the Arctic Ocean, representing an extreme in the spectrum of life-history trait variation (Nielsen 2018). Based on radiocarbon dating, females show an extraordinary lifespan estimated at 392 ± 120 y (Nielsen et al. 2016; Nielsen 2018). Longevity in this species is coupled to a very slow growth rate of <1 cm/year (Hansen 1963) and sluggishness with a cruising speed of <1 m/s (Watanabe et al. 2012). Its size and the very low temperature of its environment result in a very low mass-specific metabolic rate (Ste-Marie et al. 2020; Smith et al. 2022).

Here, we present the first chromosome-scale assembly of the GLS genome, which will be instrumental to identify the genetic adaptations responsible for this remarkable animal’s extreme longevity and peculiar traits. Our analysis suggests the presence of specific alterations in *TP53* and DNA repair.

## Results

### Chromosome-level genome assembly

To assemble the GLS genome, we used PacBio HiFi long-read sequencing. More than 50% of the initial contig assembly is covered by sequences longer than 1 Mb (**Table 1**). We performed high-throughput chromatin conformation capture (Hi-C) to assemble these sequences into scaffolds. We obtained 43 chromosome-scale scaffolds (>10Mb) accounting for a total of 6.06 Gb (94% of scaffolded nucleotides) ranging in size from 464.6 Mb for the largest- and 28.5 Mb for the smallest scaffold, respectively. A total of 12 scaffolds are longer than 200 Mb, while 17 are longer than 100 Mb (**Fig. 1a**). More than 50% of the nucleotides in our scaffold assembly are contained in sequences that are as long as the largest human chromosome (chromosome 1 with ∼248 Mb vs. our N50 of 240 Mb). With its size of 6 GB, the GLS genome is the largest shark genome assembled to date and shows the highest repeat content exceeding 70% (Table 1, compare, *e*.*g*., to (Marra et al. 2019)). Using RNA-seq and known protein sequences from other species, we identified 22,634 protein-coding genes in the genome, with 20,274 functionally annotated. The genome, annotation, and gene expression data are publicly accessible through a dedicated genome browser (https://glshark.leibniz-fli.de/). The analysis of vertebrate single-copy orthologs via BUSCO (Simao et al. 2015) indicates a genome completeness of 91.8%. Thus, despite the large size and high repeat content, the assembly is as complete as the assemblies of other high-quality shark genomes that are publicly available (**Fig. 1b**).

**Table 1.** Greenland shark genome assembly statistics.

| Total | Contigs |  |  |  | Scaffolds |  |  |  |
| --- | --- | --- | --- | --- | --- | --- | --- | --- |
| Len. | Num. | N50 | N90 | Max. | Num. | N50 | N90 | Max |
| 6.45 Gb | 13,991 | 1.1 Mb | 234 Kb | 9.2 Mb | 2,111 | 240.4 Mb | 47.6 Mb | 464.6 Mb |

**Fig 1.**
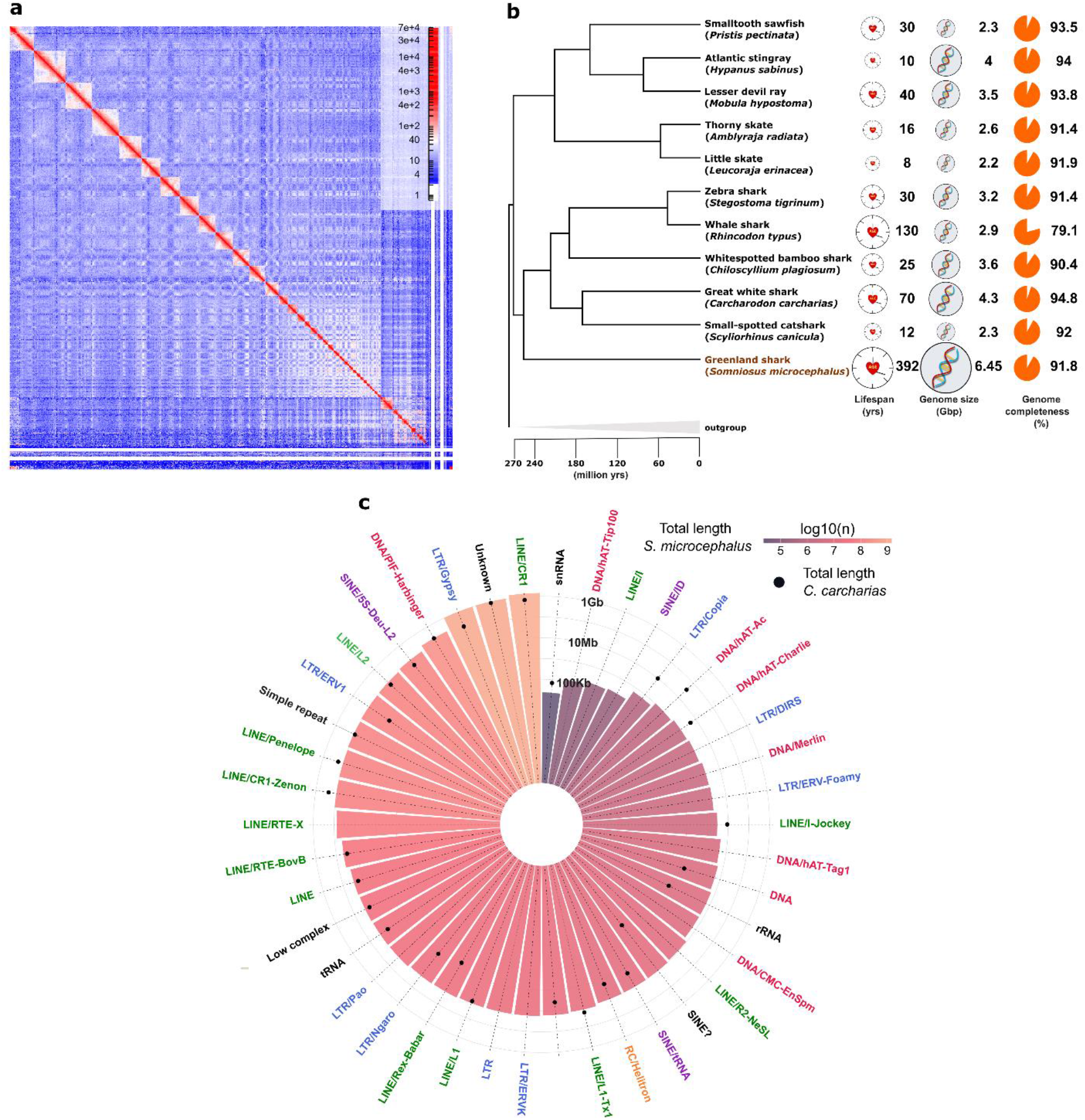
Characteristics of the Greenland shark genome assembly. a) Intensity signal heat map of HiC contacts. b) Phylogenetic tree of shark species with lifespan, genome size and genome completeness included for each taxon. Genome completeness was measured for all species by BUSCO against single-copy vertebrate genes. Divergence times were taken mainly from the timetree project (Kumar et al. 2017). Single remaining missing divergence times were taken from the following literature sources: (Marlétaz et al. 2023; Yamaguchi et al. 2023). c) Total length numbers (log10-scaled) of the most abundant transposable element families in the Greenland shark (bars). Black circles indicate the respective total length of the elements in the great white shark.

With a repeat content of 58.5%, the Great white shark (GWS, Carcharodon carcharias) was previously reported to have the highest known repeat content among sharks (Marra et al. 2019). However, the GLS genome substantially exceeds this level with a repeat content of 70.6% (**Fig. 1c**). According to our data, the increase in total content is mainly due to the expansion of retrotransposable elements, namely long interspersed nuclear elements (LINEs) and, especially, long terminal repeats (LTRs).

Taken together, we provide a highly complete, chromosome-level genome with exceptionally high repeat content for a vertebrate.

### High genomic synteny with evolutionary distant shark species

Next, we analyzed the genomic position of highly conserved vertebrate genes (BUSCO-genes, n=3,354) in the GLS and the GWS. This analysis reveals a striking synteny between the two respective genomes (**Fig. 2a**) - despite a separation between these two lineages dating about 250 million years ago (Kumar et al. 2017). For several GLS chromosome-sized scaffolds (>10Mb), we find that all BUSCO genes of any one such scaffold map exclusively to a single scaffold of the GWS genome (**Fig. 2b**). This one-to-one correspondence is observed in 19 GLS scaffolds (44% of all scaffolds). For five additional scaffolds, at least 99% of BUSCO genes mapped to the same single GWS scaffold. Thus, our data reveal very high chromosome-level synteny of 25 GLS scaffolds (60% of scaffolds) covering approximately 3 billion base pairs (46% of scaffolded nucleotides). In several instances, our synteny map indicates potential chromosomal rearrangements, *e*.*g*., involving scaffolds 11, 26, 7, and 3, as well as 29, 6, and 24. Furthermore, the map suggests potential metacentric splits of the great white shark sequences NC_04476.1 and NC_04479.1, leading to scaffolds 14 and 20 as well as 21 and 18, respectively. Three smaller scaffolds (42, 44, and 45) contained no BUSCO genes with correspondence in chromosome-scale GWS scaffolds.

**Figure 2.**
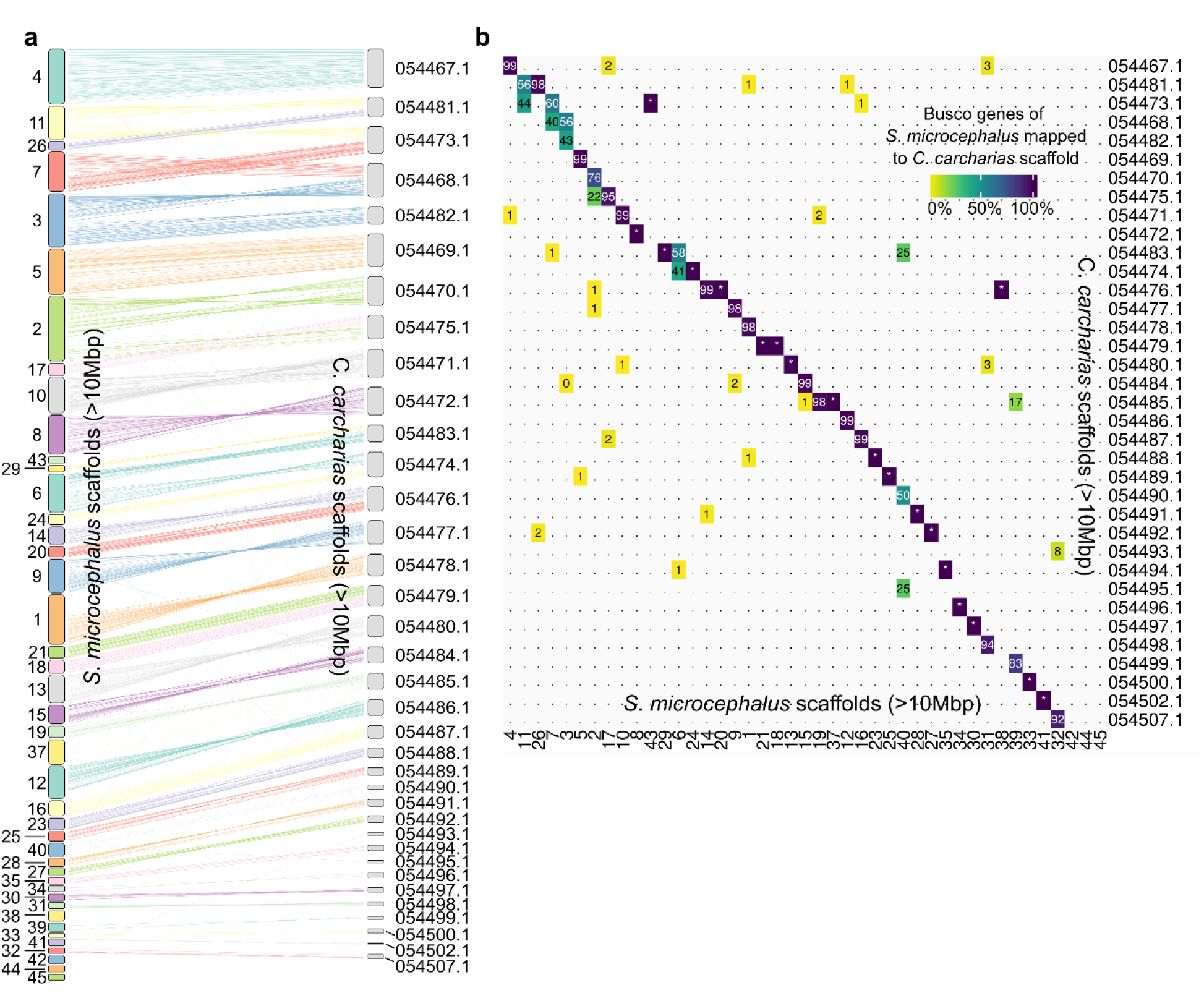
Synteny between the Greenland shark (*S. microcephalus*) and Great white shark (*C. carcharias*) genomes. a) Synteny graph of single-copy orthologous vertebrate genes found by BUSCO in both genomes. b) Heatmap quantifying for each Greenland shark scaffold (x-axis) the percentage of Busco genes mapping to the sequences of the Great White shark (y-axis). Dots indicate 0%, and stars indicate 100%.

An analysis of gene duplication events identified ancient, *i*.*e*., at least 250 million years old, duplications conserved between GLS and GWS. An extreme example of these ancient duplications is an expansion of the ferritin light chain (*FTL*) and the ferritin heavy chain (*FTH1*): we detected 155 and 179 copies of these genes, respectively. Approximately 80% of the copies are located in three larger clusters. The synteny of these clusters is identical in the GWS and GLS genomes. The largest cluster is located at the end of scaffold 30, containing 104 copies of *FTL* and 118 copies of *FTH1*. More than 85% of all *FTH1* and *FTL* copies appear to be of full length and share at least 90% of amino acids with the human orthologs. RNAseq data analysis revealed a tissue-specific expression of this cluster in the ovary (see genome browser, HiC_scaffold_30:44,903,755-48,980,971).

Overall, despite an evolutionary distance of several hundred million years between them (**Fig. 1b**), GLS and GWS show striking similarities in their genomic structure and organization.

### Specific gene duplications in the Greenland shark

Next, we sought to identify unique characteristics of the GLS genome that may underlie its phenotypic traits, particularly its longevity. Regarding gene duplications, we identified 81 genes present as single copies in all other publicly available Elasmobranch genomes (n=10) but present in multiple copies in the GLS genome. When analyzed using the STRING database, these 81 genes show significant network connectivity (p = 0.029, **Fig. 3a**), indicating that the encoded proteins cooperate in executing specific biological functions. Furthermore, this network is enriched for the gene ontology (GO) term “site of double-strand break” (GO:0035861, FDR=0.037), which is also the only GO term for which a significant overrepresentation was detected. *TP53* appears as a hub of this network, although *TP53* itself is present as a single copy in the GLS. Adding this gene manually increases the support for both network connectivity (p=0.018, **Fig. 3b**) and enrichment for double-strand break (FDR=0.002), potentially suggesting a functional link between the duplicated genes and *TP53*. Test for enrichments against the respective hallmark gene sets of the Molecular Signatures Database MSigDB (Liberzon et al. 2015) provided additional evidence for the functional relevance of the gene duplication network in DNA repair and the p53 pathway (one-sided exact Fisher-test; p=0.047 and p=0.005, respectively). Another GLS-specific gene duplication affects *NF2* coding for MERLIN, a vital tumor suppressor with known functional connections to p53 verified in mouse (Fig. 3b, (Kim et al. 2004)) and human cells (Li et al. 2017).

**Fig. 3.**
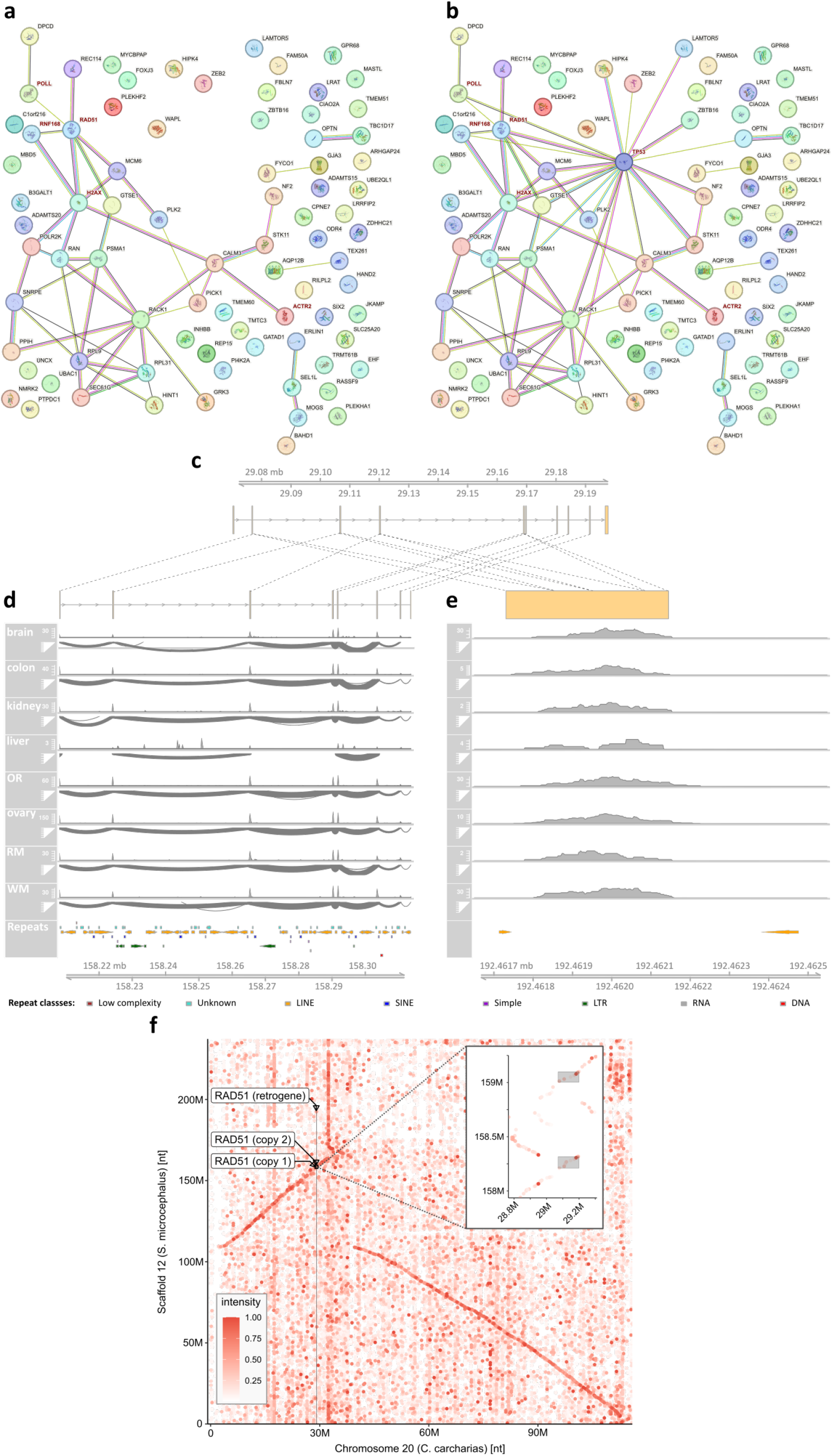
Gene duplications distinguishing the Greenland shark from other Elasmobranch species. a) Uniquely duplicated genes with potential functional connections between them (STRING database, network connectivity: p = 0.029). Red labels indicate genes contributing to the enrichment found for “site of double-strand break” (GO:0035861, FDR=0.037). b) Same as in a) with *TP53* added manually to the network. c) *RAD51* gene structure in the great white shark and two of three gene copies in the Greenland shark (d and e). Note that introns were lost in the *RAD51* copy displayed in e). At the bottom of d) and e), gene expression coverage and sashimi plots for eight tissues and repeat annotation are shown. OR – olfactory receptor; RM – red muscle; WM – white muscle. f) Alignment-based dot plot of scaffold 12 and chromosome 20 (NC_054486.1) of the Great white shark. The gene regions of the three copies of the RAD51 (indicated by arrows) show similarity with the ortholog in the Great white shark (solid vertical line). The zoom-in suggests a local duplication of a region around RAD51 in the Greenland shark. The color intensity of the dots indicates the regional sequence identity.

One of the further genes exclusively duplicated in the GLS is *RAD51*, a highly connected node in the network of duplicated genes (**Fig. 3c**), and, together with p53, its protein product plays a crucial role in DNA repair (Linke et al. 2003; Grundy et al. 2020; Laurini et al. 2020). We identified three *RAD51* copies in the GLS genome, all located on scaffold 12 in a region of approximately 30Mb. One copy of RAD51 has the same coding exons structure as in the GWS (exons 1 – 8 in the GLS; 2 – 9 in the GWS) and other Elasmobranch species displaying a high degree of conservation (**Fig. 3c,d**). Another similar copy consists of at least five exons. The third and shorter *RAD51* copy consists of a single exon (**Fig. 3e**). Of note, scaffold 12 shows a substantial level of synteny with the GWS chromosome 20 and potentially underwent an inversion of a chromosomal arm (**Fig. 3f**). Our data indicates the presence of a local duplication of a region containing *RAD51* (**Fig. 3f inset**). The single-exon copy is located about 30Mb downstream of the multi-exon copies. Interestingly, this single exon concatenates the genomic sequence of four exons of the longer copies and the corresponding four exons in the GWS and shows ubiquitous expression (exons 2 – 5, **Fig 3c**). Thus, the unique genomic organization of this *RAD51* copy may result from a retrotransposition, *i*.*e*., reverse transcription of the spliced mRNA to cDNA and subsequent integration into the genome. Several other genes exclusively duplicated in the GLS appear to result from retrotransposition. For example, *OPTN, PLEKHA1, LRAT*, and *WAPAL* have copies of only one exon matching multiple exons of the ancestral copy in the genomes of the GLS and/or other shark species. Either LINEs or LTRs flank all of these retrogenes within 1,000 base pairs.

Taken together, duplications of genes associated with DNA repair and the p53 pathway appear to distinguish the extremely long-lived GLS from other Elasmobranch species, outlining the path for additional analyses and validations.

### Positive selection and unique changes in *TP53*

We conducted a genome-wide search for positively selected protein-coding genes by analyzing the ratio between synonymous and non-synonymous substitutions to further correlate molecular and organismal evolution. So far, our analysis highlights eight genes with evidence for positive selection in the GLS, *i*.*e*., significant evolutionary changes compared to other sharks (FDR<0.1). Out of these eight positively selected genes, seven are part of pathways relevant for aging and aging-associated diseases: ZCCHC17 is a master regulator of synaptic gene expression in Alzheimer’s disease (Tomljanovic et al. 2018). ACKR3 is a chemokine receptor with pro-proliferative and anti-senescence activity (Takaya et al. 2022; Gritsina and Yu 2023). TRAPPC8, DDX42, and DESI1 are involved in autophagy, assembly of the spliceosome, and deSUMOylation, respectively (Shin et al. 2012; Zhao et al. 2017; Yang et al. 2023). A Poly (ADP-ribose) polymerase family member, PARP14, and the mitochondrial ribosomal protein MRPS35 are particularly interesting. These two genes are involved in one of two fundamental mechanisms associated with the evolution of longevity: *PARP14* in DNA repair (Nicolae et al. 2015) and *MRPS35* in mitonuclear protein balance (Houtkooper et al. 2013).

Given the role of *TP53* in longevity and cancer resistance in other large animals and its connection to the DNA repair genes duplicated in the GLS, we examined its coding sequence more closely. We detected a specific amino acid insertion in the highly conserved C-terminal region of the protein, where the GLS protein is composed of three adjacent lysines. In comparison, all other Elasmobranchs examined have only two adjacent lysines at this site (**Fig. 4a**). Notably, with the sole exception of the GLS, the two-lysine pattern has been conserved for at least ∼240 million years, which is the time when the last common ancestor of these Elasmobranchs lived (compare **Fig. 1b**). This pattern could not be detected by positive selection analysis because the probabilistic model used excludes positions that contain indels in some taxa. Structural modeling using AlphaFold3 suggests that the lysine insertion in the GLS leads to a significantly increased helix propensity and, with that, to a more extended C-terminal helix of the protein (**Fig. 4b**). Furthermore, the modeling suggests that possible interactions of the HKKEKE motif (**Fig. 4a**) with other residues of the p53 tetramer lose stability and might not be formed upon the insertion of a third lysine (K) due to the shifted position of the interaction partner (compare **Fig. 4c** to **Fig. 4d**).

**Fig. 4.**
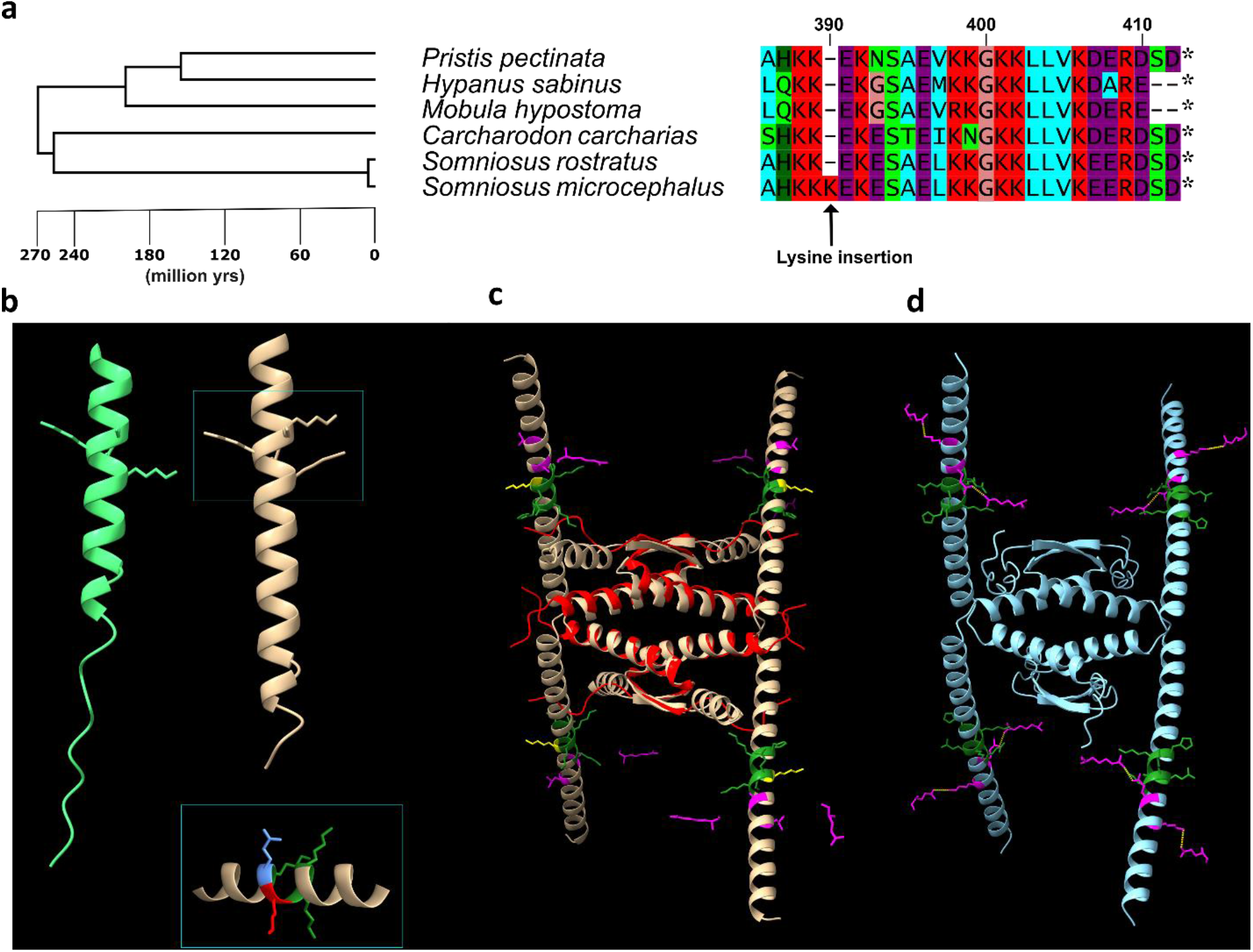
Evolutionary changes in the Greenland shark’s p53aminoacidic sequence. a) Phylogenetic tree of the protein and sequence alignment of p53’s C-terminus that contains the Greenland Shark-specific Lysine insertion. b) Side-by-side comparison of the Alphafold3 structure predictions for the C-terminal fragments of monomeric p53 of the *S. rostratus* (green, little sleeper shark) and *S. microcephalus* (Greenland shark, brown) with the latter forming a longer alpha helix. The zoom-in is showing the insertion site with its surrounding amino acids. Alignment positions K388, K389, K382 green, K390 (insertion) red, E391 in blue. c) Alphafold3 predictions for a tetrameric arrangement of p53. Superposition of the NMR solution structure of the p53 oligomerization domain (human; PDB code 1OLG; red) with the predicted model of the GLS C-terminal fragment (residues 319-409; brown). The sequence stretch H387-K-K-K390-E-K-E393 (positions by alignment, compare with a) with heavy atoms of the side chain (green) is shown with the lysine insert K390 (yellow). d) as in c) but without the K390 insertion (residues 319-408; blue). In addition, possible stabilizing interactions between the motif shown and other residues of p53 (located in different chains; E55, R310) are indicated by yellow dashes (respective residues marked in magenta).

In conclusion, our analysis shows that alterations in several aging-relevant genes, including *TP53*, distinguish the GLS from shorter-lived shark species.

### Genome Browser

To visualize and access the GLS genome and annotation, we created a genome hub for the UCSC genome browser (Nassar et al. 2023). The hub also shows GC content, repeats, BUSCO hits, and RNA-seq data of different GLS tissues mapped to the GLS genome assembly. The genome annotation and BUSCO hit tracks are searchable and can be used to find the locations and structures of genes. BLAT (Kent 2002) and *in silico* PCR (Kuhn et al. 2013) services were also set up and made available for the GLS genome assembly through the tools tab on the UCSC genome browser page. BLAT can be used to find GLS genome sequences similar to the users’ query sequence, while *in silico* PCR can be used to predict PCR product given the primer sequences and identify potential off-target sequences that may also be amplified with those primers. The hub link can be found at http://glshark.leibniz-fli.de.

Taken together, we provide an online resource for easy access to the GLS genome and its gene annotation.

## Discussion

Our study provides a chromosome-level assembly of the 6.45 Gb Greenland shark genome with an annotation of 22,634 protein-coding genes and hundreds of thousands of repetitive elements. Given the limited availability of Methuselah genomes, the Greenland shark genome provides a cornerstone for studying the genetic adaptations enabling extreme longevity.

Compared to other large deep-sea animals, we, thus far, identified fewer positively selected genes: 8 in the GLS compared to 15 in the bowhead whale, 21 in the whale shark, and 61 in the great white shark (Keane et al. 2015; Marra et al. 2019). Although few positively selected genes were found in the GLS, they may represent adaptations to multiple critical hallmarks of aging, *i*.*e*., genome instability, cellular senescence, loss of proteostasis, and disabled macroautophagy (López-Otín et al. 2023). Furthermore, we identified a network of functionally connected gene duplications in the GLS exclusively associated with double-strand break repair mechanisms and the p53 pathway, an essential regulator of DNA damage response (Williams and Schumacher 2016).

These results are in line with a large body of evidence in mammals linking double-strand break repair and longevity. For example, in elephants, the increased copy number of tumor suppressor genes, especially *TP53*, is thought to be a significant contributor to their ability to combine a considerable body size with a long lifespan (Vazquez and Lynch 2021).

Recent comparative transcriptomic studies also identified a positive correlation between the expression of DNA repair genes and the lifespan in mammals (Liu et al. 2023). For instance, the DNA repair gene XRCC6 carries a unique amino acid substitution in the Galapagos giant tortoise (Quesada et al. 2019), and duplication of DNA repair genes was detected in long-lived turtles (Glaberman et al. 2021). In addition, DNA repair genes are enriched for positive selection when comparing the longest-lived rockfish species (maximum lifespan ≥ 105 years) with the shorter-lived species of the genus (Kolora et al. 2021). At a functional level, double-strand break repair is more efficient in long-lived mammals. The somatic mutation rate scales with mammalian lifespan (Cagan et al. 2022), highlighting a pattern of evolutionary convergence to protect the genomic integrity of long-lived species. Therefore, our preliminary data may emphasize the role of DNA repair for longevity.

Furthermore, p53 itself is affected by a unique amino acid insertion in the conserved C-terminal region of the protein. This is an interesting parallel to the other very distantly related deep-sea animals, as all of these species show adaptations in key effectors of DNA repair and tumor suppression, *e*.*g*., *PARP14* / *NF2* (GLS); *ERCC1*/*ST13* (bowhead whale); *FIGL1* / *CMTM7* (whale shark); *POLD3* / *MDM4* (great white shark). Since these species can be assumed to have over a thousand times more cells than humans, it could be speculated that they are also more prone to cancer (Peto’s paradox, (Tollis et al. 2017)). Therefore, adapting the above mechanisms may have been a critical step in evading the development of malignancies.

However, an analysis of the adaptations in the four genomes of large deep-sea animals also reveals critical differences between the shorter-lived species great white shark (70 years maximum life span) and whale shark (130 years) and the even longer-lived bowhead whale (211 years) and GLS (392 years). The authors of the Great White Shark genome paper explained that they “did not find copy number variants of genes associated with genome stability”. They concluded that the positively selected genes represent “fine-tuning mechanisms related to the maintenance of genome integrity” (Marra et al. 2019). In contrast, the bowhead whale genome also contains gene duplications of individual essential DNA repair genes (Keane et al. 2015), such as *PCNA*.

Another interesting parallel concerns the shortest and longest-lived of the four species: compared to most other sharks, the great white shark (73 years) shows a substantial expansion of retrotransposon sequences (Marra et al. 2019), which is even more pronounced in the GLS (**Fig. 1c**). The expansion of retrotransposon sequences in both species contributed substantially to the increased genome sizes. In the case of the GLS, we have provided evidence that this mechanism may have contributed to the expansion of DNA repair gene sequences. Given that retrotransposons themselves are a source of double-strand breaks (Warkocki 2023), our findings suggest that the evolution of DNA repair genes and retrotransposons may be interlinked in the GLS. Specifically, retrotransposon activity may have led to the expansion of DNA repair-associated retrogenes, which, in turn, allowed for the tolerance of higher retrotransposon activity. While our observations require additional analyses and validations, it is tempting to speculate that this cycle might have been a driving force in the evolution of the GLS genome and the shark’s longevity.

## Materials and Methods

### Sample collection

Analyzed sharks were caught from 2021 - 2023 by scientific longlining from a depth between approximately 200 and 600 meters in the fjords in the vicinity of Narsaq and Qeqertoq, South Western Greenland from R/V Dana, and in Disko Bay in the area of Qeqertarsauq, North Western Greenland from R/V Porsild. All sampling was carried out following laws and regulations and with authorization from the Government of Greenland (Ministry of Fisheries, Hunting & Agriculture, permit Nr. 2020-26794 and 2023-6108).

The sharks were euthanized immediately after capture by an initial blow to the head with a bolt pistol for cattle, after which the spinal cord was transected. The tissue samples were collected as soon as possible, flash frozen in liquid nitrogen and then stored in a -80 freezer. Some samples were returned to Denmark on dry ice, and others were shipped in a freezing container.

Smaller sleeper sharks were collected in the vicinity of Genoa, Italy. One eye of 3 smaller sleeper sharks was subsequently prepared at the Aarhus AMS Centre (Department of Physics and Astronomy, Aarhus University, Denmark) by isolating the embryonic eye lens nucleus under light microscopy from concentrically arranged layers of secondary fiber cells. A 4-5 mg subsample of the innermost part of the embryonic nucleus was used for isotopic analyses with Accelerator Mass Spectrometry (AMS), as described in Nielsen et al., 2016.

### Sequencing

DNA was extracted by directly adding a digest buffer (60 mM Tris pH 8.0, 100 mM EDTA pH 8.0, 0.5% SDS) to the frozen tissue. Proteinase K was mixed into the solution until reaching a final concentration of 500 µg/ml. The mixture was kept in a water bath at 54°C until the tissue was completely dissolved. Subsequently, precipitation with phenol-chloroform was performed three times. After incubation at room temperature (3 min) and subsequent centrifugation (5 min), the aqueous phase was transferred into a fresh tube. The interphase was washed with chloroform and precipitated with 0.8 parts isopropanol. After gently swirling the mixture, it was incubated for 15 min at room temperature and centrifuged for 30 minutes. After discarding the supernatant, the pellet was washed with 70% ethanol and centrifuged again. Subsequently, the supernatant was discarded. The pellet was eluted by adding water. The mixture was placed in the refrigerator until the DNA was completely dissolved.

For Illumina sequencing, isolated DNA was quantified and quality-checked using a TapeStation 4200 Instrument in combination with Genomic DNA ScreenTape (Agilent Technologies). According to the manufacturer’s instructions, 1 µg of DNA was sheared on a Covaris M220 to a fragment size of 400 bp using the microTUBE–50 AFA Fiber Screw-Cap System. Illumina DNA libraries were prepared from 400 ng using NEBNext Ultra II DNA Library Prep Kit in combination with NEBNext Multiplex Oligos for Illumina (Unique Dual Index UMI Adaptors DNA) according to the manufacturer’s instructions (New England Biolabs). Isolated total RNA was quantified and quality checked using a TapeStation 4200 Instrument in combination with RNA ScreenTape (Agilent Technologies). Illumina RNA libraries were prepared from 800 ng using NEBNext Ultra II Directional RNA Library Prep Kit in combination with NEBNext Poly(A) mRNA Magnetic Isolation Module and NEBNext Multiplex Oligos for Illumina (Unique Dual Index UMI Adaptors RNA) according to the manufacturer’s instructions (New England Biolabs). Quantification and quality check of DNA and RNA libraries was performed using a TapeStation 4200 Instrument in combination with D5000 ScreenTape (Agilent Technologies).

Illumina DNA and RNA libraries were pooled and loaded on NovaSeq 6000 (Illumina). Sequencing was performed using a paired-end 2×101 bp configuration on S4 flow cell (Reagent Kit v1.5 200 cycles) and S2 flow cell (Reagent Kit v1.5 200 cycles). Sequencing data were converted in FASTQ format and demultiplexed using bcl2fastq software (v2.20.0.422).

HiC-libraries were prepared in part using the Arima HiC+ Kit and the Arima Library Prep Kit V2, and in part as published previously (Eremenko et al. 2021). Briefly, nuclei were isolated by homogenizing liver tissue with a Dounce Homogenizer in a fixation buffer containing 1% formaldehyde. Fixation was performed in a final volume of 10 ml of fixation buffer with moderate rotation. The reaction was quenched with 2.5 M glycine (to a final concentration of 0.2 M) for 5 minutes at room temperature with rotation. Fixed cells were washed twice, and nuclei were purified using centrifugation with a sucrose cushion. Chromatin digestion was performed by adding 200 units of DpnII restrictase per sample and incubating at 37°C overnight with constant agitation. The overhangs of the digested DNA were labeled by adding biotin-14-dATP using DNA Polymerase I, Large (Klenow) Fragment, and a mixture of other nucleotides. After pelleting the nuclei, a ligation mix (1x NEB T4 DNA ligase buffer, 1% Triton X-100, BSA, 400 U/µl T4 DNA Ligase) was added, and the DNA fragments were ligated with overnight incubation at room temperature with gentle rotation. DNA was reverse-crosslinked by incubation with Proteinase K and SDS (final concentration 2%) and then isolated using the phenol-chloroform method. DNA was sheared using a Bioruptor Pico (Diagenode) ultrasound device. MyOne Streptavidin C1 Dynabeads (Thermo Fisher Scientific) was used to pull down biotinylated fragments. Preparation for Illumina sequencing and final amplification were performed using reagents from the NEBNext Ultra II DNA Library Prep Kit for Illumina in combination with NEBNext Multiplex Oligos for Illumina (Unique Dual Index UMI Adaptors DNA) according to the manufacturer’s instructions. A mock PCR was done to finely tune the number of cycles needed for library amplification (7 cycles). Before sequencing, the size distribution and concentration of the libraries were assessed using the Agilent TapeStation System in combination with D5000 ScreenTape (Agilent Technologies). Libraries were pooled and sequenced using one NovaSeq 6000 run. System runs in SP/2x 51 cycle (paired-end)/standard-loading workflow mode. Sequence information was converted to FASTQ format using bcl2fastq v.2.20.0.422.

For PacBio sequencing, high molecular weight DNA was sheared to 7-12 kb using the MegaRuptor3 system (Speed 32, T=56min). We used the SMRTbell prep kit 3.0. Specifically, the sheared DNA was subjected to 1x MRTbell beads cleanup to reduce volume. Single-stranded DNA overhangs were removed in the first step. DNA strand breaks were repaired, and an A-overhang was added. Subsequently, non-barcoded SMRTbell adapters were ligated, followed by AmpurePB purification with two ethanol (80%) washing steps. A nuclease treatment was performed to remove residual adapters. Last, the high molecular weight fraction of the library was enriched by an AMPure PB beads size selection with a 3.1x ratio. For all runs, a total of 10 sample preps were done. All cells were sequenced on the PacBio Sequel II instrument with (10) SMRT cells using 30h video times using version 2.0 sequencing chemistry and 2h pre-extension. HiFi/CCS analysis was performed using CCS v6.3.0 with default parameters.

### Genome assembly

Contigs were assembled using hifiasm (Cheng et al. 2021) with PacBio HiFi reads as input filtered for potential contaminations using Kraken2 (Wood et al. 2019). Scaffolding was conducted to feed the contig assembly and HiC-reads into the 3D-DNA pipeline, with subsequent manual curation using a juicebox (Durand et al. 2016; Dudchenko et al. 2017). Genome completeness was measured by applying BUSCO (Simao et al. 2015) and using single-copy vertebrate genes as a reference.

### Genome annotation

Repeats were annotated using RepeatModeler and RepeatMasker (Tempel 2012; Flynn et al. 2020). Gene annotation was conducted using the braker3 pipeline (Brůna et al. 2021) with protein hints from the great white shark genome (Rhie et al. 2021) and aligned RNA-seq reads. Before this, RNA-seq reads were quality trimmed, adapter clipped, and mapped with Trimmomatic, Cutadapt, and STAR (Martin 2011; Dobin et al. 2013; Bolger et al. 2014). Functional annotation was done using eggNogMapper2 (Cantalapiedra et al. 2021). Additionally, we used BLAST (Altschul et al. 1990) to annotate unlabeled gene isoforms.

### Transcriptome assembly of *Somniosus rostratus*

RNA-seq reads were quality trimmed and adapter clipped using Trimmomatic and Cutadapt (Martin 2011; Bolger et al. 2014). Transcriptome assembly and coding sequence extraction were conducted using Trinity and TransDecoder (Haas et al. 2013).

### Comparative genomics

Analysis of transposable elements was repeated for the great white shark using RepeatModeler and RepeatMasker (Tempel 2012; Flynn et al. 2020).

Tree inference and positive selection analysis was conducted using PosiGene (Sahm et al. 2017). Coding sequences derived from our GLS genome assembly, a little sleeper shark transcriptome assembly, and publicly available coding sequences from 12 other sharks were used as input. Of the latter, ten were downloaded from NCBI (*Amblyraja radiata, Carcharodon carcharias, Chiloscyllium plagiosum, Hypanus sabinus, Leucoraja erinacea, Mobula hypostoma, Pristis pectinate, Rhincodon typus, Scyliorhinus canicula, Stegostoma tigrinum*) and two from the Squalomix webpage (*Scyliorhinus torazame, Chiloscyllium punctatum*) (Nishimura et al. 2022).

Genes duplications were analyzed by examining all gene symbols assigned more than once by the gene annotation (n=1,224). All protein sequences of all respective genes were compared against the protein sequences of ten other shark species using NCBI BLAST+. We checked for each of these ten species whether all GLS sequences associated with a specific gene symbol hit exactly one gene in the other shark species. If this was the case for each of the ten shark species for a specific gene symbol, we accepted the respective gene as exclusively duplicated in the GLS (n=81).

Dot plots and chromosome-level alignments were obtained using the D-GENIES server (Cabanettes and Klopp 2018), and visualized via ggplot using custom scripts.

### Structure prediction

Structure predictions of p53 were performed using the AlphaFold3 web server with default options (Abramson et al. 2024). For graphical representations the program ChimeraX (Pettersen et al. 2021) was employed.

### UCSC Genome Browser Hub

The make_hub.py script (Hoff 2019) was used to convert the genome assembly into 2bit file format. This script was also used to get the genome’s GC content track and to make hub descriptions and hub, genome, and track database definition file drafts. The same script was used to convert RNA-seq BAM files into bigWig files for the hub.

The genome annotation file was adjusted by ensuring correct hierarchy and attributes for gene, transcript, exon and CDS rows using gtftk (Lopez et al. 2019) and a custom bash script. The adjusted annotation was converted to USCS browser hub appropriate format using UCSC Kent Utilities gtfToGenePred, genePredToBigGenePred and bedToBigBed (Kent et al. 2010). To add BUSCO hits track, the scores in the bed file were divided by 10 to make the range of the scores compatible with the UCSC browser and bedToBigBed was used to convert the BUSCO hits bed file to USCS browser readable bigBed file format. ixIxx from USCS Kent tools was used to create indices for both the annotation and the BUSCO hits tracks to make them searchable on the hub. Repeats from RepeatMasker were converted into UCSC genome browser usable format using RepeatMasker tool rmToTrackHub.pl (Tempel 2012) followed by bedToBigBed. gfServerHuge from USCS Kent tools was used to start untranslated and translated BLAT (Kent 2002) and in silico PCR (Kuhn et al. 2013) services for the GLS genome assembly.

## Acknowledgments

The work was supported by the Leibniz Research Alliance “Resilient Ageing” with project number: LFV-2021-2-LIR to S.H. Financial support to JFS from The Danish Research Council to project “Old And Cold: Biology of the GLS”, from The Danish Center for Marine Research and the Carlsberg Foundation for a research cruise with R/V Dana to Greenland in 2021 and to the Arctic Station in Qeqertarsuaq, Greenland 2023 is gratefully acknowledged. In addition, we are thankful to EuroFleets for financially supporting the research cruise with R/V Dana in 2021 via a grant to Prof Diego Bernal, USA. We are also grateful to the crew of R/V Dana and R/V Porsild. Financial support to AC, PD, SF, and ETT from the Italian Ministry of University and Research (MUR), PRIN 2022 - SHARKAGE Grant number (CUP): E53D23007660001. The FLI is a member of the Leibniz Association and is financially supported by the Federal Government of Germany and the State of Thuringia. The publication of this article was funded by the Open Access Fund of the Leibniz Association and the Leibniz Institute on Aging – Fritz Lipmann Institute (FLI), Jena, Germany.

## Notes

### Competing Interest Statement

The authors have declared no competing interest.

